# Rethinking protein drug design with highly accurate structure prediction of anti-CRISPR proteins

**DOI:** 10.1101/2021.11.28.470242

**Authors:** Ho-min Park, Yunseol Park, Joris Vankerschaver, Arnout Van Messem, Wesley De Neve, Hyunjin Shim

**Author notes:** Corresponding Author: Hyunjin Shim. Contributions Analyses were primarily conducted by H.P., Y.P., and H.S. Specifically, AlphaFold analyses were led by H.P., genetic analyses were performed by Y.P., and structural and functional analyses were conducted by H.S. The study was conceived by H.S., and all authors contributed to writing the manuscript.

## Abstract

Protein therapeutics play an important role in controlling the functions and activities of disease-causing proteins in modern medicine. Despite protein therapeutics having several advantages over traditional small-molecule therapeutics, further development has been hindered by drug complexity and delivery issues. However, recent progress in deep learning-based protein structure prediction approaches such as AlphaFold opens new opportunities to exploit the complexity of these macro-biomolecules for highly-specialised design to inhibit, regulate or even manipulate specific disease-causing proteins. Anti-CRISPR proteins are small proteins from bacteriophages that counter-defend against the prokaryotic adaptive immunity of CRISPR-Cas systems. They are unique examples of natural protein therapeutics that have been optimized by the host-parasite evolutionary arms race to inhibit a wide variety of host proteins. Here, we show that these Anti-CRISPR proteins display diverse inhibition mechanisms through accurate structural prediction and functional analysis. We find that these phage-derived proteins are extremely distinct in structure, some of which have no homologues in the current protein structure domain. Furthermore, we find a novel family of Anti-CRISPR proteins which are structurally homologous to the recently-discovered mechanism of manipulating host proteins through enzymatic activity, rather than through direct inference. Using highly accurate structure prediction, we present a wide variety of protein-manipulating strategies of anti-CRISPR proteins for future protein drug design.

## Introduction

Proteins are macromolecules composed of amino-acid residues that perform diverse roles in biological entities, including catalysing biochemical reactions, providing cell/capsid structure, transporting molecules, replicating genetic material and responding to stimuli. It is estimated there are over 25,000 functionally distinct proteins in the human body ^1,2^, and mutations or abnormalities in these proteins may result in diseases. Thus, modern medicine has focused on harnessing target proteins to alleviate diseases, mostly through small-molecule therapeutic agents acting as competitive or noncompetitive inhibitors ^3^. However, it is estimated that only ∼10% of the human proteome can be targeted with small-molecule drugs ^3^. Since the introduction of human insulin as the first recombinant protein therapeutic in the 1980s^4,5^, protein-based therapeutics are expanding the scope of “druggable proteins”. Compared to small-molecule drugs, the major advantage of protein therapeutics is improved target specificity and reduced immunogenicity due to their proteinaceous nature ^5^. Protein therapeutics can also serve complex functions that simple chemical compounds cannot achieve, such as replacing a deficient protein or providing a protein of novel function (Fig. 1a). Furthermore, protein therapeutics can inhibit disease-related proteins that small-molecule drugs cannot target due to the lack of a cavity to bind. Currently, there are over 130 protein therapeutics commercially available and intense research efforts are ongoing to better design protein therapeutics ^5^.

**Fig. 1:**
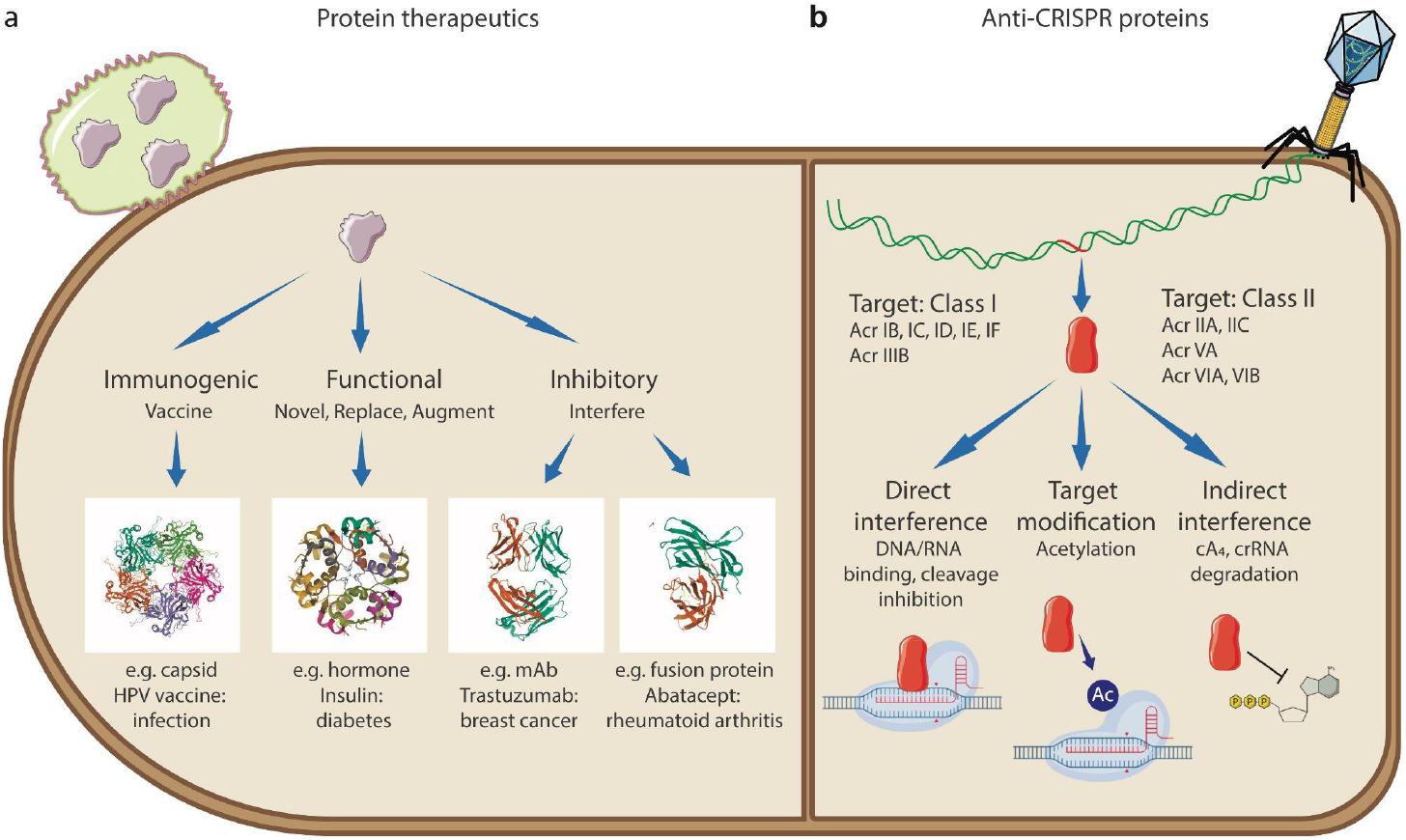
Mechanisms of protein therapeutics and anti-CRISPR proteins. **a**, Mechanism of protein therapeutics. The first group consists of prophylactic or therapeutic vaccines that induce immunity against foreign or cancer cells. A highly successful protein vaccine against human papillomavirus (HPV) combining the capsids from four pathogenic HPV strains is given as an example (PDB: 2R5K ^49^). The second group consists of protein therapeutics that provide novel functions, replace deficient or abnormal proteins, or augment existing activities. The approval of recombinant insulin in the 1980s to treat diabetes as the first abundant, inexpensive and low immunogenic therapeutic protein is given as an example (PDB: 4F8F ^50^). The third group consists of proteins that interfere with target proteins through high binding specificity. Some monoclonal antibodies use antigen recognition sites or receptor-binding domains like Trastuzumab against breast cancer cells (PDB: 6MH2 ^51^). Some fusion proteins inhibit target proteins by blocking interaction sites like Abatacept against rheumatoid arthritis (PDB: 1DQT ^51,52^) **b**, Mechanisms of anti-CRISPR proteins. Upon successful infection, phage genomes express anti-CRISPR proteins that neutralize host CRISPR-Cas immunity. Anti-CRISPR proteins target various types of both Class I and Class II CRISPR-Cas systems and the inhibitory mechanisms are highly diverse, including direct interference of DNA/RNA binding and cleavage of Cas complexes, enzymatic inhibition of the active site by acetylation, and nuclease activity of degrading the signaling molecule (cyclic nucleotide cA_4_).

In this study, we present a group of naturally-occurring protein therapeutics called anti-CRISPR (Acr) proteins, as a prominent example of how small proteins are used by invading bacteriophages (phages) in nature to control host proteins. Phages are the most abundant and diverse biological entities in the biosphere (estimated ^31^ phages) that infect and replicate within host prokaryotes (such as bacteria or archaea) ^6^. High selective pressures between these parasites and hosts drive dynamic coevolution of genomic and proteomic mechanisms and systems ^7–9^. In particular, the evolutionary arms race between phages and prokaryotes has resulted in a vast arsenal of immune systems, including the prokaryotic adaptive immune system known as CRISPR-Cas ^10,11^. CRISPR-Cas systems are defence mechanisms against phages (and other mobilomes) through a complex of RNA-guided Cas proteins (Fig. 1b). Remarkably, prokaryotic genomes with CRISPR-Cas systems can acquire short fragments of foreign genetic sequences in their CRISPR arrays, which serve as RNA templates to recognize and cleave invading phages through the nuclease complex of Cas proteins ^12^. Since the successful application of CRISPR-Cas systems as genome-editing tools ^13,14^, there was a burst in the discovery of diverse CRISPR-Cas systems ^15^, followed by the discovery of Acr proteins that neutralize the activity of this prokaryotic adaptive immune system ^16^ (Fig. 1b). A family of Acr proteins was first identified in the CRISPR-Cas-inactivating prophages of *Pseudomonas* genomes that disable Type I-F and Type I-E CRISPR-Cas systems ^16,17^. A number of Acr proteins inhibiting type II CRISPR-Cas systems have since been applied as regulators of gene-editing activities ^18^. Acr protein families are known to have short sequences (<100 amino acids) with no common genetic features, and interact directly with Cas proteins to inhibit target DNA binding, DNA cleavage, CRISPR RNA loading and protein-complex formation ^18,19^. A recent study reveals AcrVA5 proteins to inactivate Cas12a of Type V CRISPR-Cas systems enzymatically by acetylation of the active site, with structural homology to an acetyltransferase protein ^20^.

In this study, we conduct a comprehensive analysis on the key characteristics of Acr proteins viewed from the perspective of naturally-occurring protein therapeutics that effectively inhibit host protein functions. Motivated by the observation that these Acr proteins are genetically diverse, we examine the protein structure of these diverse proteins using AlphaFold ^21^. AlphaFold is a state-of-the-art deep learning-based approach that performs protein structure prediction, which takes a protein sequence as an input to predict its 3-D protein structure through an iterative exchange of information between its genetic representation and its structural representation. The recent release of AlphaFold, the winner of CASP14 which achieves highly accurate protein structure predictions ^21,22^, is revolutionary for the field of life sciences and medicine, and is expected to accelerate critical research in a large number of fields ranging from structural biology to drug discovery. In this study, we first assess the performance of AlphaFold in predicting the 3-D structures of Acr proteins based on similarity metrics against their experimentally reconstructed 3-D macromolecular structures. Using this performance as a basis, we further examine the Acr proteins without experimental structures with AlphaFold, to predict the structural diversity of these genetically distinct proteins that are natural inhibitory proteins against prokaryotic CRISPR-Cas systems. We use AlphaFold-predicted structures of Acr proteins to infer a range of inhibition mechanisms through homology search and functional analysis, to demonstrate how bacteriophages exploit diverse strategies to manipulate host immune systems, with the long-term goal of providing an unique opportunity to learn from the evolution-optimized inhibitor proteins for future protein drug design.

## Results

### AlphaFold prediction of anti-CRISPR protein structures

Acr protein datasets were acquired from various viral and prokaryotic genomes, including *Pseudomonas phage, Pseudomonas aeruginosa* and *Escherichia coli* ^23^, which were categorised into three sets: verified Acr proteins with experimental structure (Set A), verified Acr proteins without experimental structure (Set B), and putative Acr proteins with experimental structure (Set C) (Supplementary Table 1). From the AlphaFold-predicted structures of each set (Fig. 2a), we compared the prediction performance between CASP14 (52 AlphaFold evaluation results from the CASP14 competition), Set A and Set C (Fig. 2b and 2c), based on TM-scores and relative Z-errors (see Methods for details) against the true experimental structures (Supplementary Table 2), where Set B without experimental structures was excluded. According to the TM-score, CASP14 had a higher median than Set A and Set C (0.925 vs. 0.895; 0.843, respectively), but Set A had the highest mean as compared to Set C and CASP14 (0.896 vs. 0.868; 0.882, respectively). Furthermore, Set A and Set C had significantly smaller standard deviations than CASP14 (0.095; 0.092, respectively, vs. 0.120), indicating that, according to the TM-score, their predictions are more accurate. According to the relative Z-error, CASP14 recorded a lower median than Set A and Set C (0.201 vs. 0.217; 0.230, respectively), but Set A recorded the lowest mean as compared to CASP14 and Set C (0.211 vs. 0.237; 0.259, respectively). Similarly to the TM-score, Set A and Set C recorded smaller standard deviations than CASP14 (0.128; 0.129, respectively, vs. 0.141). As can be seen in the boxplots (Fig. 2b and 2c), there was no significant difference in prediction performance, validating that AlphaFold predicts 3-D structures of Acr proteins as accurately as the CASP14 dataset.

**Fig. 2:**
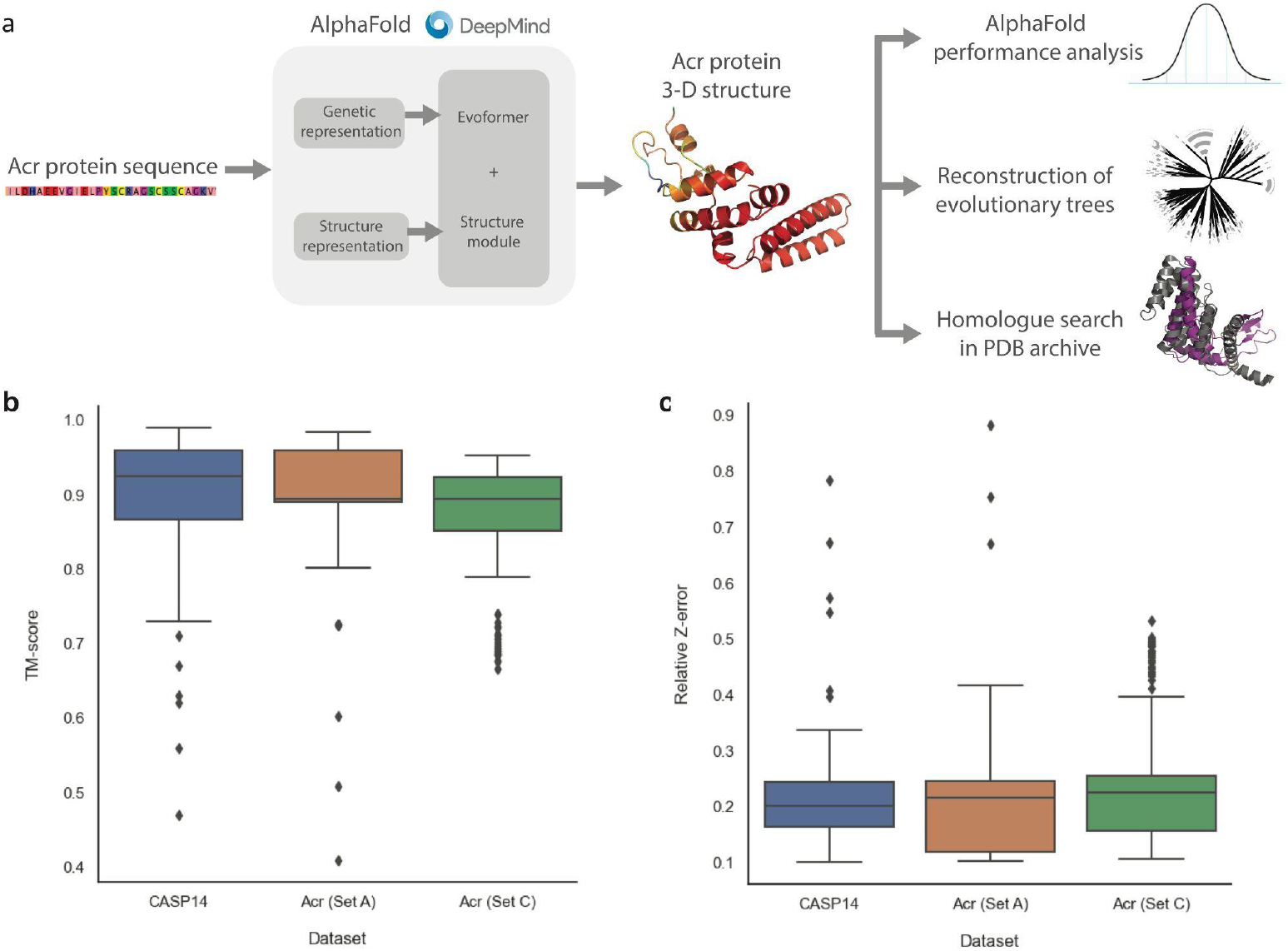
Performance analysis of AlphaFold on anti-CRISPR proteins in comparison to the CASP14 dataset. **a**, Overall workflow to analyze the 3-D macromolecular structures of Acr protein sequences predicted with AlphaFold. **b**, The performance of AlphaFold on the Acr protein datasets in comparison to the CASP14 dataset using TM-scores. The closer the TM-score is to 1, the more similar the predicted structure is to its true experimental structure. **c**, The performance of AlphaFold on the Acr protein datasets in comparison to the CASP14 dataset using relative Z-errors. The closer the relative Z-error is to 0, the more similar the predicted structure is to its true experimental structure. (Set A: Verified Acr proteins with experimental structures, Set C: Putative Acr proteins with experimental structures).

### Evolutionary trees of Anti-CRISPR proteins

Acr proteins are known to be genetically diverse; this raises an intriguing question about the origin and evolution of Acr proteins. We reconstructed evolutionary trees of the verified Acr proteins (Set A + Set B; *n*=207), using sequence-based and structure-based methods (see Methods for details). As expected, the phylogenetic tree built using genetic sequences (Fig. 3a) shows high levels of variation, while forming consistent clades with the high bootstrap support. Analysis of the clades reveals some degrees of clustering by the Acr family at the shallower nodes; however, this clustering is mostly due to the near-identical protein sequences. For instance, many of the AcrIIA proteins were derived from AcrIIA1 and AcrIIA2, driven by the technological interest to regulate CRISPR-Cas gene-editing activities in different cell types ^18^. Otherwise, the phylogenetic tree shows absence of clustering by other biological features such as taxonomy and inhibition mechanism.

**Fig. 3:**
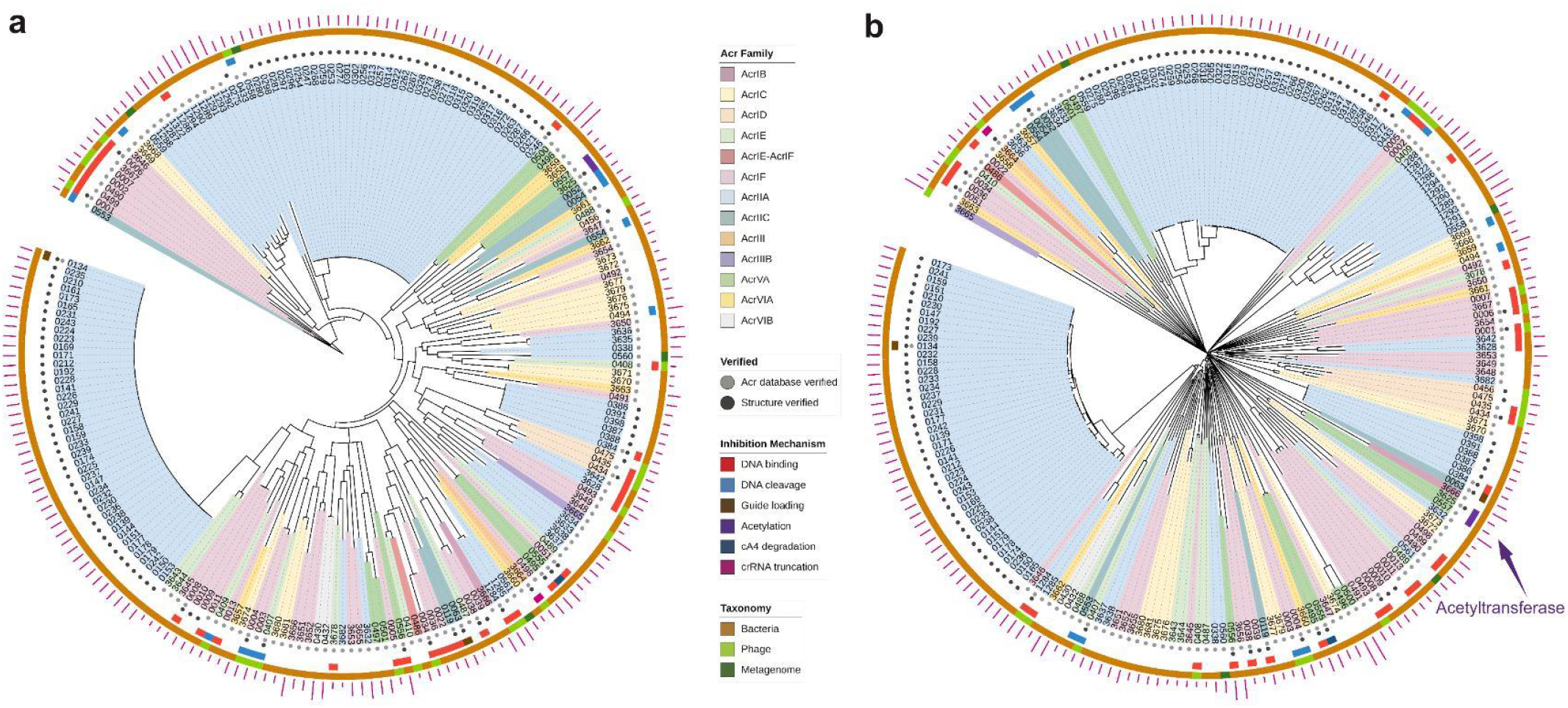
Evolutionary trees of anti-CRISPR proteins. **a**, The phylogenetic tree of anti-CRISPR proteins reconstructed using sequence-based methods (Set A + Set B; *n*=207). **b**, The structural tree of anti-CRISPR proteins reconstructed using structure-based methods (Set A + Set B; *n*=207). The clades of the two evolutionary trees were coloured by the Acr Family. The inner to outer rings display the Acr verification status, structural verification status, inhibition mechanism and source organism taxonomy. The outer magenta bars represent the genetic sequence length of each protein.

Given the low sequence similarity among the Acr proteins, we built a structure-based tree using AlphaFold-predicted structures, which includes 98 Acr proteins without experimental structures (Fig. 3b). The structural tree shows an even higher level of diversity in the Acr proteins than the phylogenetic tree. In the structural tree, the Acr proteins share no common ancestor and display deep branches, consistent with earlier observations of how evolutionary pressure drives immunity-related mechanisms of hosts and parasites to coevolve rapidly ^8,9^. The structural tree also shows some degrees of clustering by the Acr family, but the clusters do not always coincide between the two evolutionary trees. From the visual analysis of the protein structures, the branches of the structural tree are placed randomly in terms of structural forms and the functions are only related at the clade level (Supplementary Fig. 1).

It is evident that the sequence-based and structure-based trees capture different evolutionary relationships between the Acr proteins. 3-D structures of homologous proteins were previously shown to be better conserved than their corresponding genetic sequences, particularly when the sequence similarity is below 30% ^24^. From the multiple sequence alignment, no site of the Acr proteins is conserved at 30% and only very few sites are conserved at 15% (Supplementary Fig. 2). We calculated the congruence among distance matrices of the sequence-based and structure-based trees to be very low according to the measure of congruence (Kendall’s coefficient of concordance, W = 6.58e-01), confirming the correlation between these two types of evolutionary trees is poor among highly divergent proteins ^24^.

### Structural homology to predicted anti-CRISPR structures

Characterised Acr proteins use diverse strategies to interfere directly with CRISPR-Cas systems, including inhibiting DNA binding, DNA cleavage, guide loading and ribonuclease activity ^18^. To investigate the relation between 3-D protein structures and inhibitory functions, we identified homologous structures to the AlphaFold-predicted Acr proteins using structure-based distance measures ^25^ (see Methods for details).

First, we used a subset of the Acr dataset with experimental structures (Set A) to successfully validate the closest structural homologue to each AlphaFold-predicted Acr protein matched with its true experimental structure (Supplementary Table 3).

Second, we analyzed the Acr proteins without experimental structures (Set B) by matching the AlphaFold-predicted structures to the closest homologues from the Protein Data Bank archive ^25^ (Supplementary Table 4). We first validated that Acr proteins retrieved their neighbours in the same clade as the closest homologue, when their neighbouring proteins had experimental structures in the Protein Data Bank. For example, the Acr proteins labelled ‘0434’ and ‘0435’ are in the same clade, and the closest homologue of the Acr protein ‘0435’ matched with the experimental structure of the Acr protein ‘0434’. Functional analysis of the closest homologue to each Acr protein reveals a wide variety of protein functions, including polymerase, ligase, nuclease, regulation, and transport (Fig. 4c). We acknowledge that some of the closest homologues have low structural similarity (Z-score below 4); however, it is intriguing that these Acr proteins have no close structural homologues in the Protein Data Bank. Some Acr proteins from the families AcrIC6, AcrVIA2, AcrIIA19, and AcrIF8 (Fig. 4a) have no structural homologues (significance threshold for similarity: Z-score = 2).

**Fig. 4:**
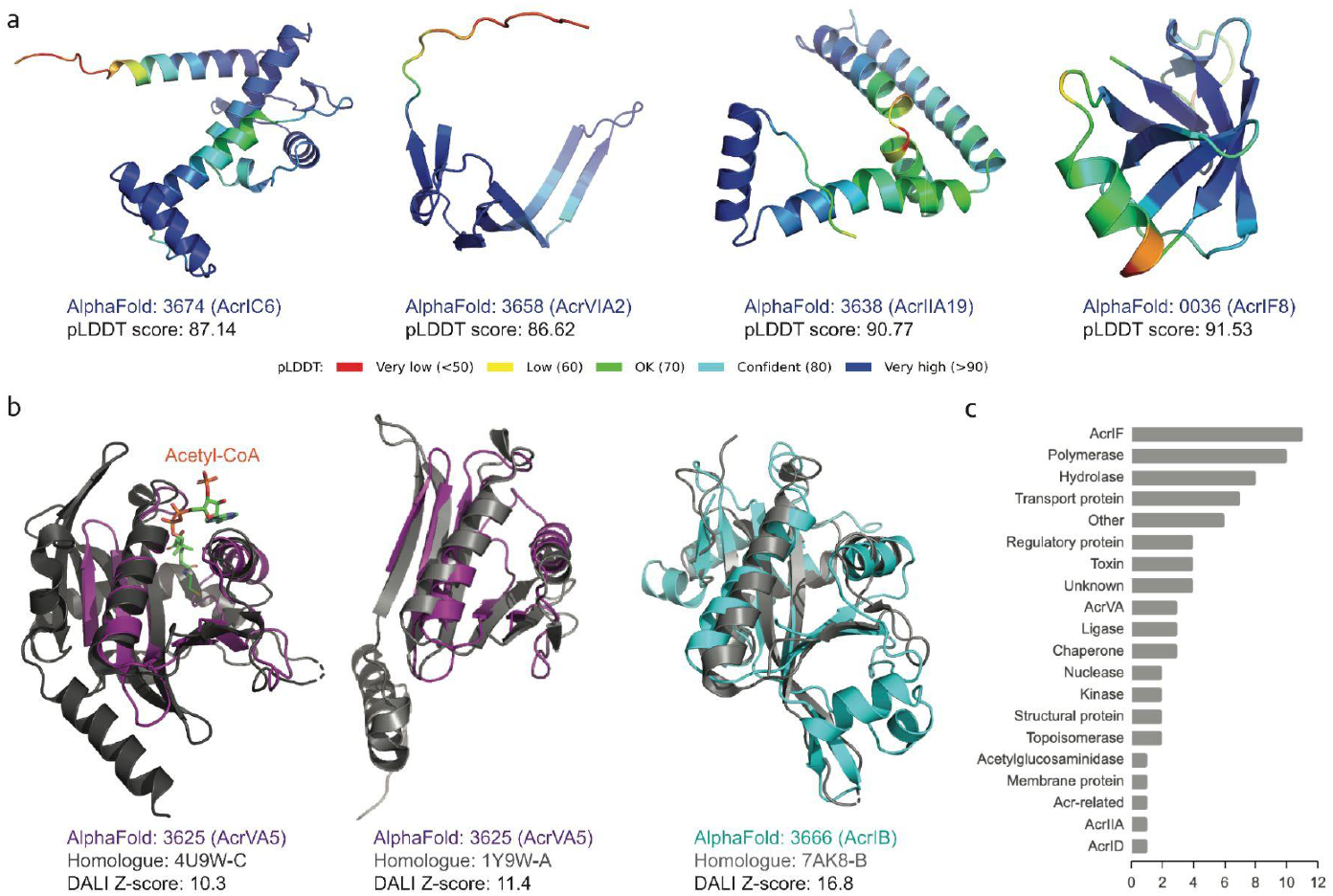
AlphaFold-predicted 3-D structures of outlier anti-CRISPR proteins. **a**, The AlphaFold-predicted structures of the Acr proteins without homologues in the Protein Data Bank archive (Dali Z-score < 2). The 3-D protein structures are coloured according to the b-factor spectrum in PyMol, with a per-residue estimate of the AlphaFold confidence on a scale from 0 - 100 (high pLDDT accuracy in blue, low pLDDT accuracy in red). **b**, Superimposition of the AlphaFold-predicted structures of Acr proteins and their closest structural homologues retrieved from the Protein Data Bank archive. All the closest structural homologues in grey have functional annotations related to acetyltransferase. The homologue to the Acr protein labelled ‘3625’ has a cofactor (acetyl-CoA) bound, revealing the functionally critical site of the enzyme. **c**, Functional analysis of the closest homologues to the AlphaFold-predicted Acr proteins without experimental structures (Set B). Only the functional annotations above the significance threshold of Dali Z-score (>4) were included.

Third, we cross-examined the Acr proteins whose inhibitory mechanism was experimentally characterised to verify that the homologues retrieved were functionally related (indicated as the middle ring in Fig. 3b). The homologue functions of this subset were related to a wide variety of functional domains (Supplementary Table 4). For instance, the two Acr proteins in the same clade (labelled ‘3628’ and ‘3642’) were characterised to inhibit the DNA-binding of Cas proteins ^26^ and their structural homologues have functional domains of ligase. A few closest homologues with functions related to acetyltransferase drew particular attention, as a recent biochemical study revealed an unprecedented mechanism of inhibiting CRISPR-Cas systems through enzymatic activity rather than through direct interaction ^20^. According to this study, the closest structural homologue to this Acr protein (labelled ‘3625’) was found to be N-Alpha-Acetyltransferase from *Homo sapiens* (4U9W-C) (Fig. 4b), despite their low sequence similarity. We found another homologue (1Y9W-A) with a better similarity score to the AlphaFold-predicted structure of this Acr protein, that had the functional annotation of acetyltransferase from *Bacillus cereus* (Table 1). In addition, we found a number of uncharacterised Acr proteins in the same clade of the structural tree (between ‘0430’ and ‘3681’) related to Acetylglucosaminidase from various Acr families, including AcrIC, AcrIE, AcrIF, AcrIIA, and AcrVIB. Intriguingly, several proteins have homologues with the functional annotations of nuclease activity, which is reminiscent of the newly-discovered mechanism of nuclease activity against crRNAs and CRISPR-Cas signalling molecules ^27,28^.

**Table 1.** Closest homologue to the AlphaFold-predicted structure of Acr proteins with acetyltransferase annotations.

| Acr ID | Family | Type | Length | Homologue | Dali Z-score | Annotation | %ID structure | %ID sequence |
| --- | --- | --- | --- | --- | --- | --- | --- | --- |
| 3625 | AcrVA5 | V-A | 92 | 4U9W-C | 10.3 | N-Alpha-Acetyltransferase | 14 | 6.8 |
| 3625 | AcrVA5 | V-A | 92 | 1Y9W-A | 11.3 | Acetyltransferase | 22 | 16.8 |
| 3666 | AcrIB | I-B | 193 | 7AK8-B | 16.8 | Acetyltransferase | 21 | 17.6 |

%ID structure: percentage identity in structure
%ID sequence: percentage identity in sequence

### New anti-CRISPR family of acetylation inhibition

Previously described Acr proteins of the Acr family VA5 disable Type V Cas12a by acetylation, which leads to a complete loss of the DNA-cleavage activity ^20^. We found that this AcrVA5 protein (labelled ‘3625’) structurally aligned closer to *Bacillus cereus* acetyltransferase than the previous structural homology of *Homo sapiens* acetyltransferase (Fig. 4b). We further identified other Acr proteins that are related to this AcrVA5 protein on the evolutionary trees of the Acr proteins (Fig. 3b). Notably, there are two Acr proteins in the same clade as the AcrVA5 protein on the structural tree, one of which was lacking experimentally validated structure or function. Analysis of its function using the structural homologues reveals that this Acr protein (labelled ‘3666’) is related to acetyltransferase. Interestingly, this Acr protein belongs to a different Acr family (AcrIB) than the previously identified AcrVA5. Its superimposition with the closest structural homologue reveals a similar structural alignment at the functionally critical site of the acetyltransferase where acetyl-CoA binds (Fig. 4b). The sequence identity of the AcrIB protein to its closest homologue was found to be 17.6%, while the structural identity between these two proteins in 3-D was found to be higher at 21% (Table 1 and Supplementary Fig. 3). On the phylogenetic tree, this AcrIB protein is not placed close to the other two Acr proteins of acetyltransferase function (labelled ‘3625’ and ‘0557’) (Fig. 3a), demonstrating that these two types of Acr proteins have close structural similarity but not genetic similarity. This finding suggests that for proteins with low sequence similarity, structure-based trees cluster proteins with most similar biochemical functional properties better than sequence-based trees ^24^. Using the structural tree, we discovered a new family of Acr proteins belonging to AcrIB that had a strong structural homology to acetyltransferase from a different organism (gram-negative bacteria *Salmonella enterica*), whereas the previously characterised AcrVA5 matched to acetyltransferase from gram-positive bacteria *Bacillus cereus*.

## Discussion

We show that the 3-D structures of Acr proteins predicted with AlphaFold achieve high accuracy. The structural tree reconstructed from these AlphaFold-predicted structures display more diversity of Acr proteins with no common evolutionary origin as compared to the phylogenetic tree. On the structural tree, the Acr proteins form small clades by the unique structural similarity, which are also related by the inhibition mechanism. The functional annotations of the Acr protein homologues are extremely diverse, including a wide range of enzymatic and regulatory activities from different organisms. Most characterised Acr proteins inhibit host CRISPR-Cas systems by direct interference; we show that this category of Acr proteins displays various functional annotations and unique structural forms in the multiple branches of the structural tree.

In particular, we found a number of Acr proteins with homologue annotations related to acetylation. A recent discovery of Acr proteins that manipulate CRISPR-Cas systems through enzymatic activities demonstrates extensive phage defence mechanisms driven by the intense host-parasite arms race ^20,27^. Through the AlphaFold-predicted structural analysis, we found a novel family of Acr proteins (AcrIB) from the genome of a human pathogen (*Leptotrichia buccalis* C-1013-b) that shows more structural similarity to acetyltransferase than the previously characterised AcrVA protein. Intriguingly, other Acr proteins on the multiple branches of the structural tree have homologues related to different types of acetyltransferase enzymes from heterologous species. Acr proteins could evolve independently from various host genomes and mobile genetic elements, exploiting a vast inventory of protein structures as the basis for their counter-defence advantage ^6,9,29^.

More broadly, Acr proteins are exceptional examples of coevolution dynamics optimizing the phage genomes to manipulate host systems and maximize survival. Since phages can only replicate within host cells and are void of metabolic capacity to synthesize small molecules, their counter-defence machinery against the sophisticated and extensive prokaryotic anti-phage systems is protein-based. Nonetheless, phages are the most abundant organisms in the biosphere ^30^, and their successful protein-based viral arsenals such as Acr proteins provide an important insight on how to expand the potential of protein therapeutics against disease-causing proteins. Specifically, we could get inspiration from these phage-derived protein structures that resemble segments of related or target proteins, to design highly-specialised protein inhibitors with diverse protein-manipulating strategies, including indirect interference through enzymatic activities ^31^. The high biodegradability issue of protein therapeutics has partially been solved by the recent success of mRNA vaccine delivery using lipid nanoparticles ^32^, making low risk protein therapeutics ever more attractive to the industry. From the AlphaFold-predicted structures, we accelerated the structural and functional analysis of Acr proteins whose experimental 3-D structures are yet to be resolved. Given the number of Acr proteins without homologues in the current protein structure domain, we wonder whether there is a vast repertoire of unexplored protein structural configurations that can be exploited for protein drug design.

## Methods

### Curation of anti-CRISPR datasets

The Anti-CRISPR dataset contains 446 Acr proteins (210 verified, 236 putative) that inhibit a wide range of CRISPR-Cas systems including I, II, III and VI from the Anti-CRISPRdb ^23^. The term ‘verified’ indicates the protein was validated as CRISPR-Cas-inactivating Acr (either by the database or other published papers), while ‘putative’ indicates the protein was predicted to be Acr without sufficient experimental support. The Anti-CRISPR dataset was further curated into Set A, Set B, and Set C for AlphaFold, according to the availability of experimentally reconstructed 3-D macromolecular structures (hereby referred to as “experimental structures”) (Supplementary Table 1). Each protein is annotated with Acr family, type of inhibited CRISPR-Cas systems, NCBI accession, genetic sequence, source organism, taxonomy, and inhibitory mechanism when available (Supplementary Tables 3 and 4).

### Prediction of anti-CRISPR protein structure with AlphaFold

We predicted 3-D protein structures of each set with AlphaFold, using the Acr protein sequence as the input to AlphaFold (Fig. 2a). AlphaFold creates genetic and structure representations by comparing the protein sequence with several pre-installed databases. Those representations are used as input to five prediction models to generate five candidate 3-D structures. The result with the highest per-residue confidence score (pLDDT: per residue estimate of confidence on a scale from 0 to 100 ^33^) among the five results is determined as the final structure and saved in a protein data bank (PDB) format (Supplementary Fig. 4). For the Acr datasets, we used the PDB archive until 31/12/2012 as templates in AlphaFold to exclude the true experimental structures of the Acr proteins. The details of the experiments related to the hardware specification and to the processor performance are given in the Supplementary material (Supplementary Table 5 and Supplementary Table 6, respectively).

### Comparison of AlphaFold performance on anti-CRISPR against CASP14

To validate the performance of AlphaFold on predicting Acr structures, we benchmarked the CASP14 dataset against Set A and Set C of Acr proteins with the corresponding true experimental structures available. We excluded predicted structure and experimental structure pairs for which the TM-score and/or the Z-score were too low. Finally, 52 pairs of CASP14, 99 pairs of Set A, and 207 pairs of Set C were used for the comparison study. We used the TM-score ^34,35^ and Dali Z-score ^36^ as similarity measures between the predicted and the experimental structures. Unlike traditional metrics (e.g. root-mean-square deviation), the TM-score is length-independent and more sensitive to the global similarity than to the local variation. The Dali Z-score is the sum of the equivalent residue-wise intermolecular distances among two proteins, and does not have a fixed upper bound ^36^. We then used the following relative Z-error to calculate the relative difference:

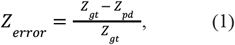

where *Z*_*gt*_ is the self Dali Z-score between experimental structure and itself, and *Z*_*pd*_ is the Dali Z-score between experimental structure and predicted structure. We obtained the *Z*_*pd*_ and TM-score for the CASP14 set of AlphaFold from the CASP14 assessment scores ^37^, whereas *Z*_*gt*_ of CASP14 and *Z*_*gt*_ and *Z*_*pd*_ of Set A and Set C were calculated using DaliLite.v5 ^36^. Finally, a protein structure comparison and clustering tool called MaxCluster ^38^ was used to calculate the TM-scores of Set A and Set C. Both distance metrics have values between 0 and 1, with 1 as the best score for TM-scores and 0 as the best score for relative Z-errors.

### Reconstruction of evolutionary trees of anti-CRISPR proteins

We reconstructed the evolutionary trees of the Anti-CRISPR dataset (Set A and Set B) using sequence-based and structure-based inference. The sequence-based tree of the Acr proteins was built by aligning the amino acid sequences using a multiple alignment program, MAFFT (version 7.471, -auto option) ^39^. The multiple sequence alignment of the Acr proteins was then visualized using Jalview (version 2.11.1.3) with a conservation visibility of 15% (Supplementary Fig. 2) ^40^. Subsequently, a phylogenetic tree of the Acr proteins was built with IQ-Tree using ModelFinder (-auto option) to find the best-fit model among the supported range of protein substitution models ^41,42^ (Supplementary Table 7). Using the best-fit substitution model, 1,000 ultrafast bootstrap replicates were run to check bootstrap support of the reconstructed tree topology ^43^.

The structure-based tree of the Acr proteins was built by calculating the similarity matrix between the Dali Z-scores of the AlphaFold-predicted structures and its corresponding experimental structures. We used the Dali server ^36^ for generating structural trees from hierarchical clustering of the similarity matrix. The structural tree of the Acr proteins was generated from distance matrices, where the pseudo-distance between two structures Q and T was defined as ^44^:

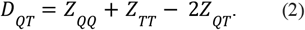

The hierarchical clustering of the similarity matrix was outputted as a Newick formatted dendrogram. The phylogenetic tree and the structural tree of the Acr proteins were visualized with iTOL (version 4) and iTOL annotation editor ^45,46^ with the following labels: Acr Family, Taxonomy, and Inhibition Mechanism.

### Congruence among distance matrices of sequence-based and structure-based trees

We measured the congruence among distance matrices of the reconstructed trees from the sequence-based and structure-based methods using Kendall’s coefficient of concordance, W, which ranges from 0 (no congruence) to 1 (complete congruence) ^47^. First, we computed the cophenetic value of pairwise distances between the terminals from a phylogenetic tree using its branch lengths with the function `cophenetic.phylo` from ape-package (version 5.0) ^48^. Then, we used the function `CADM.global` to calculate the coefficient of concordance among the distance matrices of the sequence-based and structure-based trees of the Acr proteins through a permutation test.

### Visualization of protein structure superimposition

For functional analysis, AlphaFold-predicted structures with functional annotations of interest were superimposed with their structural homologues using PyMol (version 2.5.2) to visualize the overlap in structure of the functionally active sites. The inhibitory mechanism of Acr proteins without experimental structure was inferred through examining functional annotations of the structural homologues to the AlphaFold-predicted structure, with the significance threshold of Z-score > 4.

## Data Availability

All the input protein sequences are available in Anti-CRISPRdb (https://guolab.whu.edu.cn/anti-CRISPRdb) as well as our project GitHub page (https://github.com/powersimmani/ACR_alphafold). The 3-D structures from the Protein Data Bank were used as ground truth for calculating Dali Z-scores and TM-scores; and superimposing structures (downloaded on 13/10/2021). All AlphaFold-predicted 3-D structures for Acr proteins are available on our github project page, including 3-D images. TM-scores and Dali Z-scores of AlphaFold; ground truths of CASP14 were acquired from the CASP (Critical Assessment of Structure Prediction) competition (https://predictioncenter.org/casp14).

## Code Availability

Protein structures were predicted with AlphaFold, available under an open-source license at https://github.com/deepmind/alphafold. For protein structure similarity metrics, we used MaxCluster (http://www.sbg.bio.ic.ac.uk/~maxcluster/index.html) for TM-score and DaliLite.v5 (http://ekhidna2.biocenter.helsinki.fi/dali/README.v5.html) for Dali Z-score. For MSA, we used MAFFT.v7 (https://mafft.cbrc.jp/alignment/server) and Jalview.v2 (https://www.jalview.org) for visualization. For phylogenetic tree reconstruction, we used IQ-Tree (http://www.iqtree.org) with ModelFinder and UFBoot options. For structural tree reconstruction, we used Dali server (http://ekhidna2.biocenter.helsinki.fi/dali) for building dendrograms. 3-D Structure visualizations were created in Pymol v.2.5.2 (https://pymol.org) and Py3DMol v.1.7.0 (https://pypi.org/project/py3Dmol) with Jupyter v.1.0.0 (https://jupyter.org). For data analysis, Python .3.6.4 (https://www.python.org), NumPy v.1.17.5 (https://github.com/numpy/numpy), SciPy v.1.1.0 (https://www.scipy.org), seaborn v.0.9.0 (https://github.com/mwaskom/seaborn), Matplotlib v.3.3.4 (https://github.com/matplotlib/matplotlib), pandas v.0.22.0 (https://github.com/pandas-dev/pandas) were used.

## Acknowledgements (optional)

The research and development activities described in this study were funded by Ghent University Global Campus (GUGC), Incheon, Korea.

## Supplementary Information

Supplementary Information will be provided upon request.

